# Persistent Invasion Risk: Modeling the near-Current and Future Distribution of Pterygoplichthys disjunctivus across the Philippine Archipelago

**DOI:** 10.64898/2026.05.10.724170

**Authors:** Jean-Matthew B. Bate, Almyt Poblete, Nikki Heherson A. Dagamac

## Abstract

Philippine freshwater ecosystems are considered one of the most diverse ecosystems harboring numerous fish species. However, in the Philippines, these ecosystems are threatened by invasive species that potentially disrupt ecological balance. In this study, we focused on the vermiculated sailfin catfish Pterygoplichthys disjunctivus, an invasive aquarium species reported in several Philippine aquatic ecosystems. Despite its documented spread, its potential range under a rapidly changing climate remains poorly understood for the country. Hence, in this study, we utilized the MaxEnt model to predict its near-current and future habitat suitability in the Philippines. Using 11 reported occurrences, our model showed high predictive accuracy (AUC = 0.882± .034, TSS = 0.7394 ± 0.154, SEDI = 0.971 ± 0.019). Across the current and future scenarios, slope was the primary contributor (78.7% – 81.3%), followed by BIO 10 or the mean temperature of the warmest quarter(18% – 27.8%), and flow accumulation (0% – 5.2%). However, for the SSP126 scenario, BIO10 is projected to triple by 2050 (18 – 48%). Current projections identify high-risk regions, particularly central Luzon (Laguna de Bay and Lake Taal), the Cagayan River Valley, and portions of eastern Mindanao (Agusan Marsh and Lake Mainit). Sankey transition analysis confirms a high habitat stability rate (>73%) for high-suitability pixels in both SSPs, indicating persistent invasion risk. Overall, our study provides a framework for invasive species management and contributes to the conservation of Philippine aquatic ecosystems.

## INTRODUCTION

The Philippines harbors diverse freshwater ecosystems, accounting over 374 freshwater fish species, 64 of which are introduced (Jamandre, 2023). The country’s archipelagic nature supports a vast number of freshwater ecosystems encompassing over 406,328 hectares of socio-ecologically and economically important inland waters, including eight of the RAMSAR-recognized wetlands of international importance (Sespeñe et al., 2016; Guerrero, 2022). Although known for sustaining rich freshwater biodiversity, the Philippines faces several challenges of environmental and anthropogenic pressures, one of which is the alarming spread of invasive species (Palmer et al., 2015). Among all threats to Philippine inland waters, the introduction of potentially invasive species poses the second-highest threat to biodiversity loss, next to habitat destruction (Guerrero, 2022). These invasions carry heavy ecological and financial costs; For example, the persistent increase in the catch of the Midas cichlid *Amphilophus citrinellus* in Lake Taal suggests continued spread, thereby threatening to displace native species such as the *Sardinella tawilis* through overpopulation, affecting the lake’s ecological balance and livelihood of the community (Mutia et al. 2025).

However, addressing these biological invasions requires more than just monitoring; it underscores the need for predictive understanding of the species’ niche requirements to predict potential range expansion, especially in the Philippines where inland waters are abundant and interconnected. Maximum entropy (MaxEnt) modelling has emerged as a robust tool for this case, particularly when dealing with presence-only data, typical of emerging invasions. While this modeling method has been successful in predicting the potential expansion range of invasive fishes in Asia (Mamun et al. 2018; Vythalingam et al. 2022; Marr et al. 2024) its application is timely in the Philippines to address the invasive vermiculated sailfin catfish *Pterygoplichthys disjunctivus*. A famous aquarium fish that is native to South America which already has expanded its distribution to countries in North America (Nico et al. 2006; Grana, 2007; Rueda-Jasso et al. 2013) and Asia (Suresh et al. 2019; Prakash et al. 2022; Limbu et al. 2024) including the Philippines (Chavez et al. 2006; Bate et al. 2025).

The species was first reported in the country in late 1990s - presumed through aquarium release and aquaculture escapes- and was formally documented in Laguna de Bay by Chavez et al. (2006). Subsequent reports indicate that the species is significantly more widespread than previously thought, even reaching far north of Luzon island, and as far as south as the Agusan Marsh in Mindanao island. The species possess highly adaptive traits that allow it to tolerate a wide range of environmental parameters (Suresh et al. 2019; Dien et al. 2023). These include high fecundity, tough armor-like scales, and specialized physiological traits that allow it to persist in hypoxic and polluted aquatic environments. Furthermore, its burrowing behavior has been reported to cause significant soil erosion and water turbidity, physically altering the water body it invades (Santos and Velez-Gavilan, 2024). Ultimately, once established, the species is known to displace native species and disrupt ecological balance (Bostami et al. 2025)

Hence, our study aims to predict the near-current and future scenarios of *P. disjunctivus* in the Philippines using MaxEnt modelling to identify its habitat suitability under changing climate scenarios. By identifying high risk areas, we provide a framework for targeted management and conservation efforts of Philippine freshwater ecosystems against the introduction of a highly invasive species.

## METHODOLOGY

### Species occurrences

A total of 12 *P. disjunctivus* occurrences were determined from Bate et al.’s (2025) updated locality record of *P. disjunctivus* (Figure 1). To prevent model overfitting due to spatially clustered occurrences, we used the *spThin* package (Aiello□Lammens et al. 2015) of R software, which has delimited the points within a 10 km radius, resulting in 11 total points for the model.

**Figure 1.**
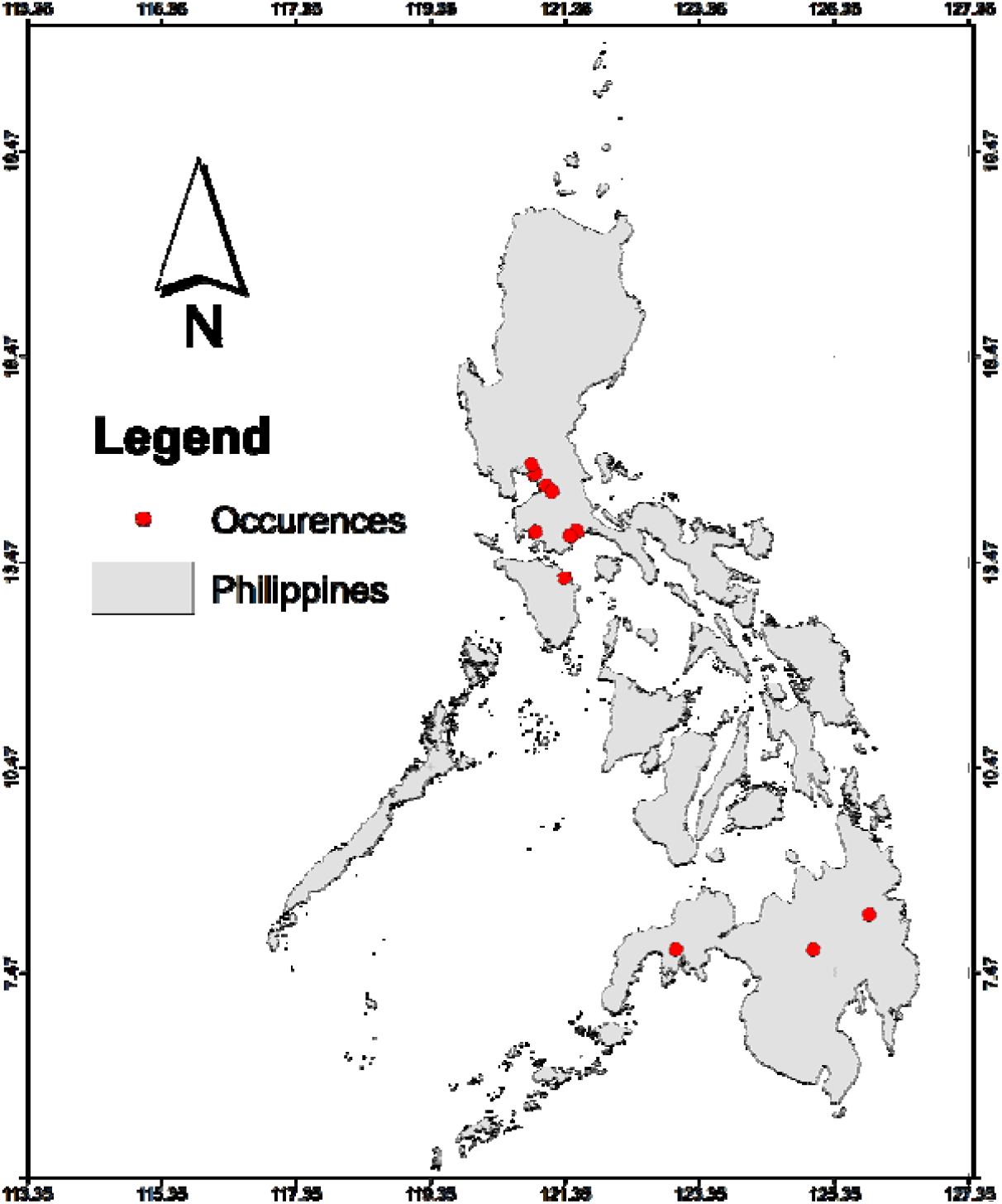
Map of the Philippines with the 12 *P. disjunctivus* occurrences.

### Environmental predictor selection

To determine the current and future distribution of the invasive sailfin catfish in the Philippines, we used the dynamic climatic layers in combination with the static topographical and hydrological layers. We collated the historical-current (1970-2000), 2030 and 2050 standard 19 BioClim layers, representing Shared Socioeconomic Pathway (SSP) 126 and 585 from www.worldclim.org. SSP126 refers to the optimistic sustainable scenario where society is geared towards carbon mitigation. Meanwhile, the SSP585 refers to the pessimistic scenario, simulating fossil-fueled development scenarios without mitigating global warming conditions. The topographic and hydrological variables (elevation, slope, flow directions, and flow accumulation) were pre-processed in ArcGIS from the digital elevation model (DEM) of the Philippines (CGIAR-CSI, 2016). All environmental variables were resampled to a spatial resolution of 30 arc-seconds (∼1 km^2^) and reprojected to the WGS84 (EPSG:4326) geographic coordinate system using R studio to ensure spatial consistency.

To address multicollinearity among layers, we conducted Pearson correlation using *Terra* (Hijmans et al. 2022) and *caret* (Kuhn, 2008) package of R studio. Variables with a correlation coefficient of >0.7 were considered redundant. Ultimately, an optimal number of three environmental predictors (Pearson et al. 2006; Breiner et al. 2015) were chosen (BIO10, flow accumulation, and slope) to predict the habitat suitability of *P. disjunctivus:* The climatic variable BIO10 (the mean temperature during the warmest quarter) represents thermal stress brought by the rapidly changing climate scenarios, which is also known to facilitate invasion of this species (Carlson, 2025; Gilles et al. 2025). Slope was selected due to the sedentary nature of the sailfin catfish observed at the bottom of water bodies (Santos and Velez-Gavilan, 2024). Higher slopes typically correlate with faster moving waters, while lower slopes represent flat, stagnant waters. As a substrate-associated grazer, these species are highly adapted to depositional water areas where organic detritus accumulate (Hubilla et al. 2007). Lastly, flow accumulation was included to represent hydrological connectivity that has sufficient water and established river systems needed by a large-bodied invasive fish (Santos and Velez-Gavilan, 2024).

### MaxEnt modelling, validation, and evaluation

MaxEnt v3.4.4 (Philips et al. 2017) was used for species distribution modelling using presence-only data of *P. disjunctivus* in the Philippines. We used bootstrap subsampling approach with 10 replications, which is optimal for few occurrences such as this case. The occurrence data were partitioned into 75% for model training and 25% for independent testing (omission error and AUC evaluation). Furthermore, we ran the model with default parameters –10,000 maximum background sites, iteration of 500, using the auto-feature for the parameter regularization, and using “1” as the regularization multiplier. Jackknife was used to identify the importance of each predictor variable. The output format was set to Cloglog.

To assess the current scenario’s model accuracy and performance, we determined three (3) indices, allowing robust model assessment compared to relying on single evaluation method, which increases potential bias (Shabani et al. 2018). First, we utilized the built-in statistics computation of MaxEnt to determine the (1) area under the curve (AUC) where values <0.7 indicate poor model performance, 0.7 – 0.9 indicates moderate performance, and values >0.9 indicate good model performance (Pearce and Ferrier, 2000). Next, the (2) true skill statistics (TSS) were identified by selecting MaxSSS (maximum sum of sensitivity and specificity) as the threshold (Liu et al. 2016) and calculated using the formula:

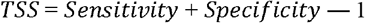

TSS values ranges from 0.2 – 0.5 indicating poor performance, 0.6–0.8 is useful, and values > 0.8 is deemed as excellent model performance (Gama et al. 2017; Li et al. 2023). Lastly, we calculated the Symmetric Extremal Dependence Index (SEDI), which has recently emerged as an optimal model skill evaluation test for presence-background analyses that are appropriate for limited sample sizes (Bouam et al. 2022; Wunderlich et al. 2019). Unlike TSS which is sensitive to species prevalence, SEDI is prevalence-independent and non-degenerate reflecting true environmental suitability for limited sample sizes. It is calculated as:

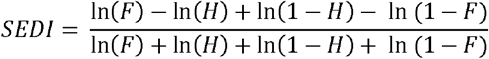

Where F is false alarm rate (1 – Specificity) and H is hit rate (Sensitivity).

## RESULTS

### MaxEnt model performance and validation

The AUC, TSS, and SEDI values (mean±SD) from the 10 bootstrap subsampling replicates resulted in 0.882± .034, 0.7394 ± 0.1537, and 0.971 ± 0.019, respectively. The minimum and maximum values for each model are: 0.601 and 0.993, 0.514 and 0.993, and 0.940 and 0.999, respectively, showing high consistency across multiple replicates. The averages of the validation metrics indicates that the MaxEnt model for the current scenario has indeed reliable predictive performance (Figure 2).

**Figure 2.**
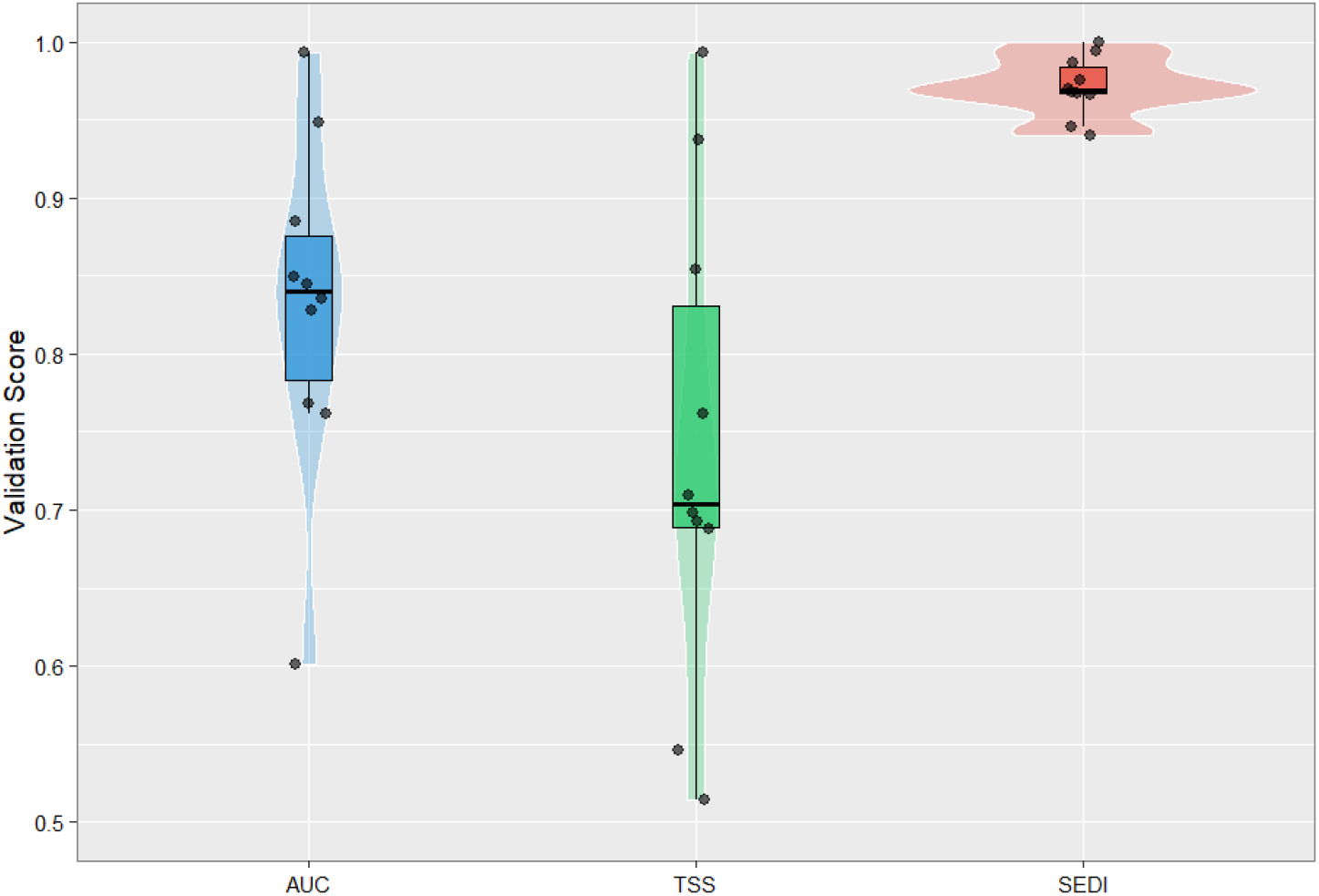
Model evaluation metrics (AUC, TSS, and SEDI) for *P. disjunctivus* distribution modeling.

### Predictor importance

Across the three scenarios, slope emerged as the primary predictor of *P. disjunctivus* habitat suitability (78.7% – 81.3%). This is followed by BIO10 ranging from 18% – 27.8%, and flow accumulation with 0% – 5.2%. However, yearly transitions of predictor contribution are different for the two pathways (Figure 3) Under the SSP126 scenario, BIO10 drastically increases from 18 % – 48% by 2050. In contrast, under the SSP585 scenario, the dominance of slope intensifies, reducing the contribution of BIO10 and ultimately masking flow accumulation to zero by 2050.

**Figure 3.**
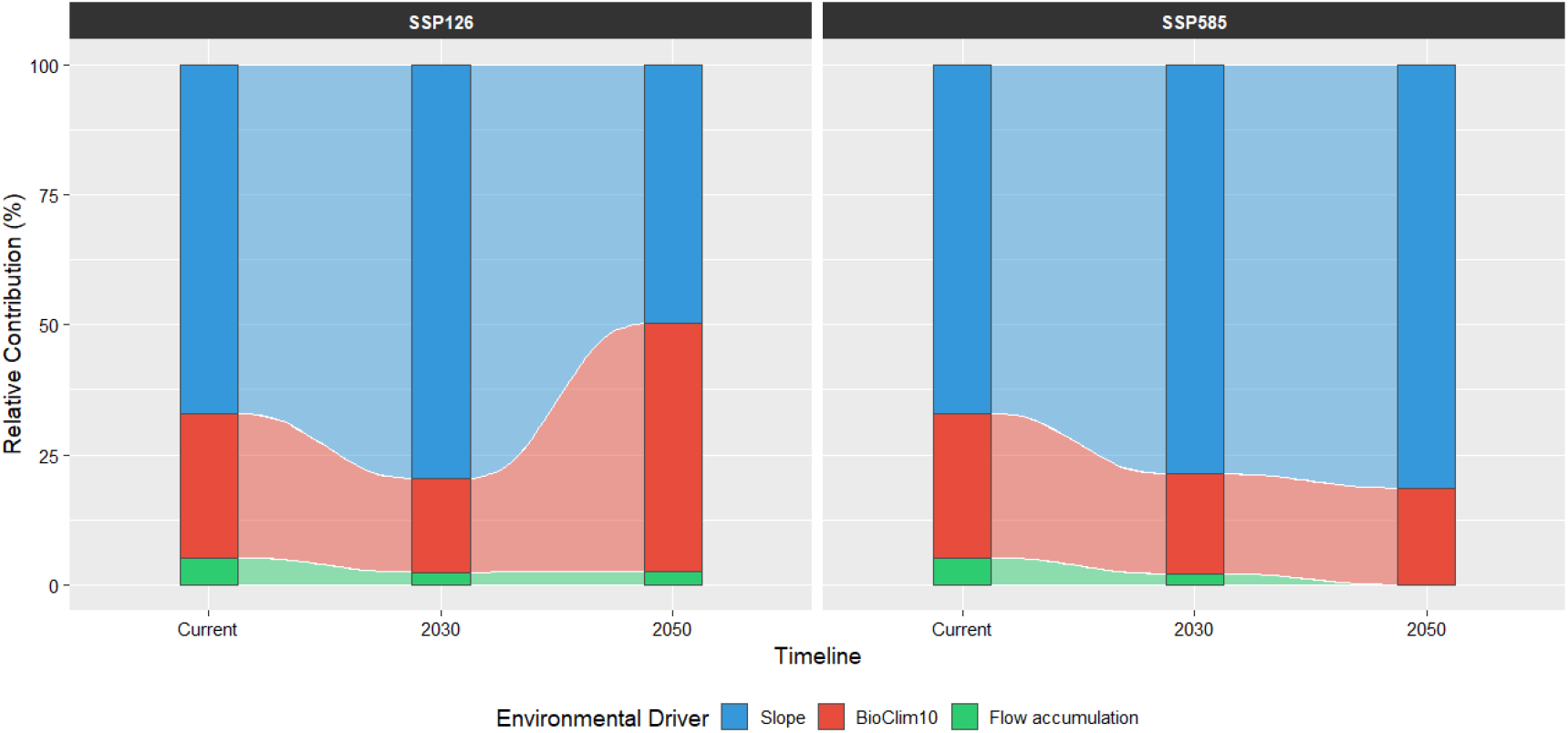
Predictor contribution transition for *P. disjunctivus* under two shared socioeconomic pathways (SSP126 and SSP2050).

### Current and future habitat suitability of P. disjunctivus in the Philippines

The current distribution map of the species in the country indicates high environmental favorability in major freshwater drainage systems across the Philippines (Figure 4). Highly suitable regions are most prominent in central Luzon basins, including Laguna de Bay and Lake Taal, both of which are major lakes, as well as Cagayan River Valley in the northern portion of Luzon. High suitable areas appear fragmented in the Visayas region, particularly the western portion, and the lowland coastal areas of Negros occidental and Northern Leyte. Within Mindanao Island, highly suitable areas are prominent in the Southern portion and in Agusan province basins which include the protected Agusan Marsh and Lake Mainit. Medium suitable areas aggregate in Southeastern Luzon, eastern Visayas, and central Mindanao including Lanao Lake, which is a major lake in the province.

**Figure 4.**
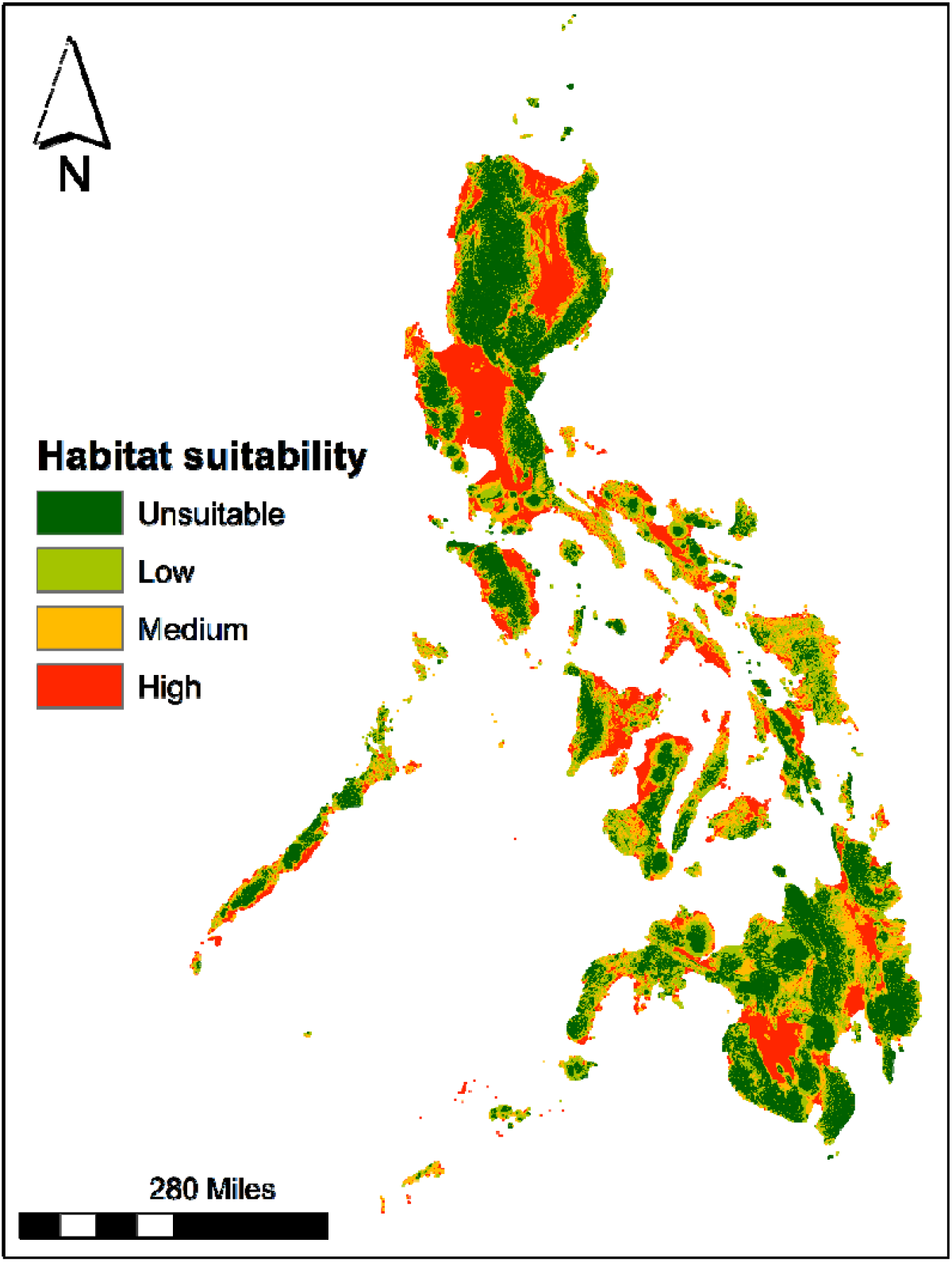
Current habitat suitability of *P. disjunctivus* in the Philippines.

Comparative analysis of spatial projection between SSP126 and SSP585 shows a high degree of spatial similarity of the future distribution of *P. disjunctivus* from 2030 to 2050 (Figure 5). Similar to that of the current scenario, the same high suitability basins from central Luzon and Mindanao Island are observed. The only notable change is the shift of high suitability to medium suitability in eastern Mindanao from SSP126 2030 to 2050. This is further confirmed by our Sankey transition analysis (Figure 6), indicating the same pixels remain relatively stable across different socio-economic pathways up to 2050. Under the SSP126 scenario, 77.6% of pixels classified as “high suitability” remained in the same class by 2050, while 96.8% of unsuitable pixels were retained. In contrast, pixels classified as low and medium suitability were more sensitive to change with lower stability rate of 67.3% and 62.7%, respectively.

**Figure 5.**
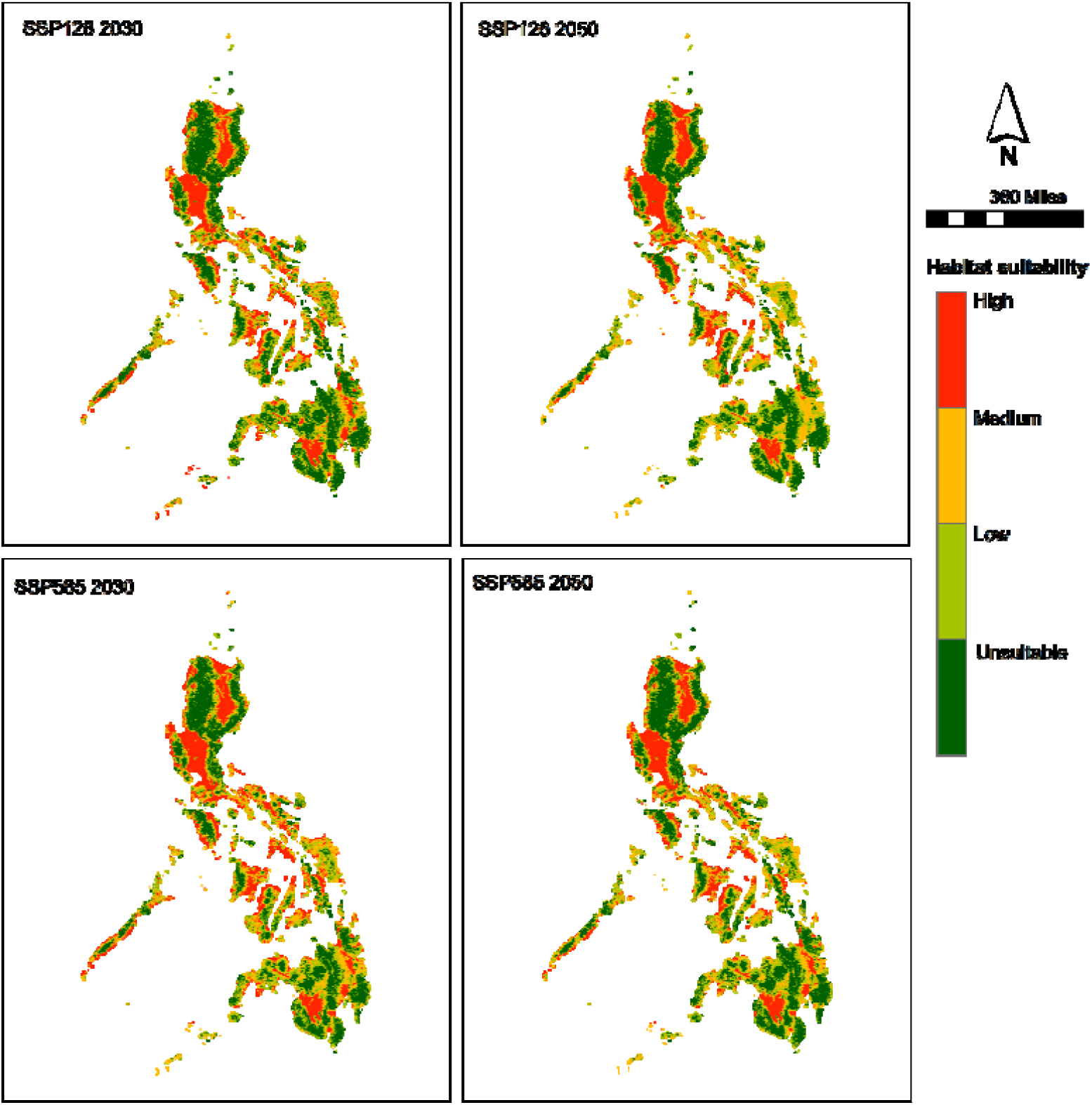
Future distribution range prediction of *P. disjunctivus* using MaxEnt model.

**Figure 6.**
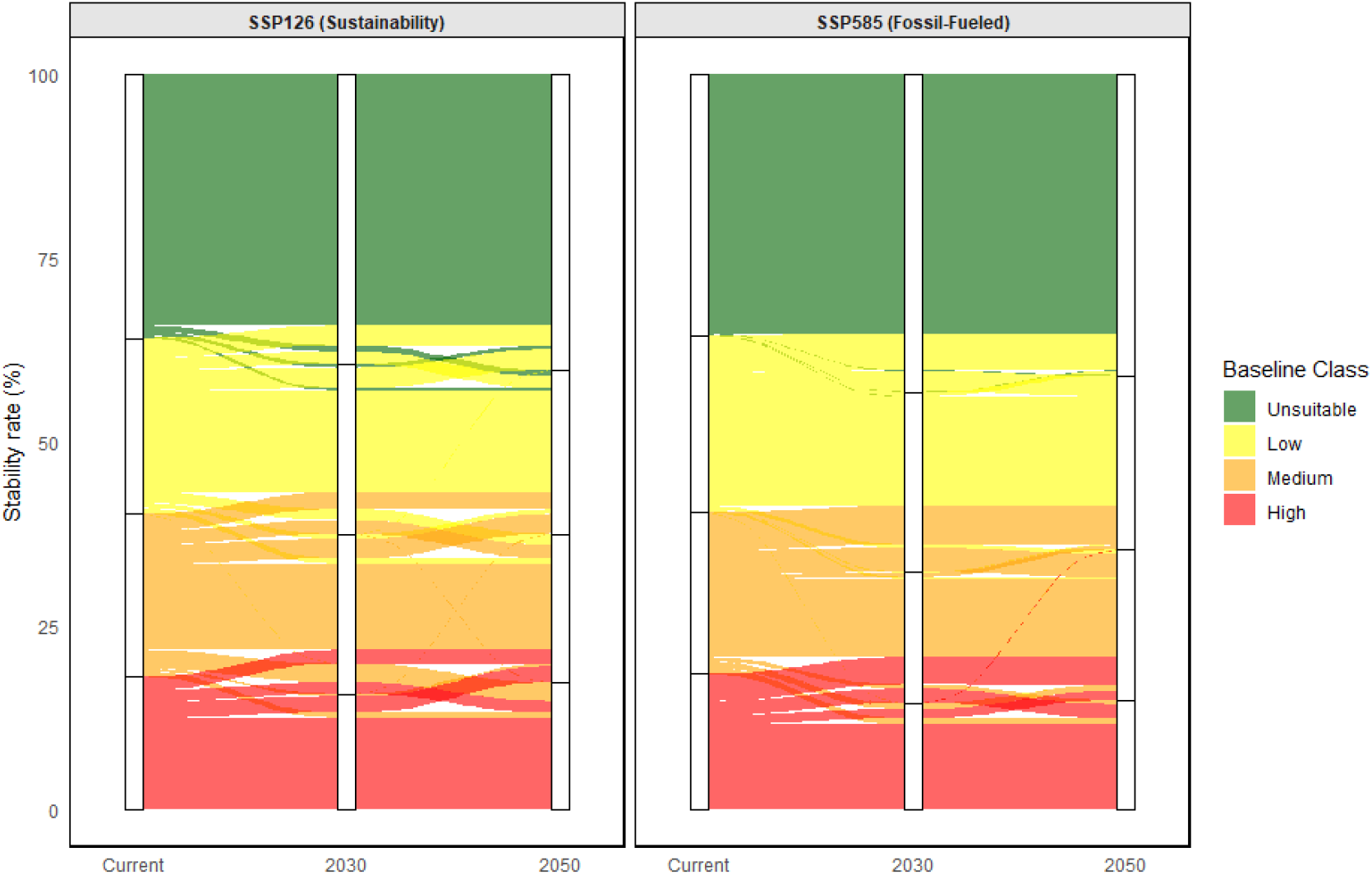
Sankey diagram showing habitat suitability transitions (represented as percent of pixels retained) for *P. disjunctivus* from current conditions to 2050.

Under the SSP585 scenario, the stability of habitat suitability was more pronounced. The pixels classified as unsuitable remained 99.5% stable, while highly suitable pixels maintained a stability rate of 73.1%. The low and medium suitability pixels, similar to that of SSP126, represented zones of higher suitability change.

## DISCUSSION

*Pterygoplichthys disjunctivus* is one of most recognized invasive fish species globally, including the Philippines (Orfinger and Goodding, 2018). Despite this, monitoring through predictive surveillance modelling in the country is still overlooked, which may lead to future introductions and potential harm to its megadiverse ecosystems. This study provides a robust MaxEnt model of *P. disjunctivus* distribution, that was thoroughly evaluated using three independent validation tests through a country-level prediction of the Philippines.

### Current and future distribution of P. disjunctivus

The invasive sailfin catfish are predicted to be widely distributed in central and northern Luzon basins, fragmented portions of Visayas region, southwest and northeast Mindanao, particularly covering lowland flat regions (slope) with warm temperatures (BIO10) and high volumes of water (Flow accumulation) (see supplementary files). Alarmingly, this region comprises several protected lakes that are both economically and ecologically important. These water bodies are inhabited by native and endemic species (*Arius manillensis, Cestraeus goldiei, Gobiopterus lacustris, Leiopotherapon plumbeus, Sardinella tawilis*) which may compete with the sailfin catfish if introduced (Aquino et al. 2011; Escaño et al. 2022; Gilles et al. 2023). This includes Laguna de Bay in Laguna province, Lake Taal in Batangas province, Lake Paoay in the Ilocos region, Cagayan River Valley in the Northern portion, and Lake Naujan in Southern Luzon. In the Visayas region, Lake Danao in Cebu, which is inhabited by the native *Anguilla marmorata* and *Clarias batrachus* (Romero et al. 2023), is at high risk. While in Mindanao Island, the Agusan Marsh and Lake Mainit are at high risk.

Based on our future projections, the *P. disjunctivus* display high environmental resilience across both sustainable and fossil-fueled climate pathways. The similarity between current and future projections suggests that the core invasion areas in the country are environmentally suitable through 2050. This is further confirmed by our Sankey transition analysis, showing high stability rate (% of pixel retained) throughout 2030 to 2050. The high stability rate may have been caused by the dominance of topographic variable (slope) as a predictor compared to the climate variable, which is expected to change. As a generalist, the species is known to thrive in a wide range of thermal conditions (Orfinger and Goodding, 2018), even thriving in tropical waters (Hossain et al. 2018; Stolbunov et al. 2021). Consequently, increase in mean temperature may not act as a physiological barrier for *P. disjunctivus* distribution. Based on our model, climate is secondary to depositional low-velocity environments. This suggests that as long as the hydro-geographic structure of the basins remain unchanged, the species will be able to persist regardless of socio-economic pathway.

### Potential introduction impacts and implications for freshwater ecosystem management

If left unmanaged, the species may adversely affect the ecological and economic integrity of important freshwater systems causing severe degradation. Particularly in areas where *P. disjunctivus* introduction has been reported such as in Agusan Marsh, the species have been linked to significant diet overlaps with native fishes and increased turbidity and erosion (Hubilla et al. 2007). Beyond ecological effects, the species are also a threat to food security and livelihood; Their rapid reproduction (Gibbs et al. 2017; Hossain et al. 2018) leads to overpopulation, decreasing the palatable fish catch (Chavez et al. 2006) and causing financial strains to fishermen by destroying gill nets (Chavez et al. 2006; Bate et al. 2025). In other countries, the potential impacts of *P. disjunctivus* include change in pond structure due to its burrowing behavior, alter nutrient dynamics, and may ultimately bring novel parasites to the aquatic ecosystem (Capps and Flecker, 2013a, 2013b; Nitta and Nagasawa, 2016; Hussan et al. 2022).

Despite existence of national legislation against introduction of non-native species in the Philippines (Republic Act 9147, the Wildlife Resources Conservation and Protection Act), accidental introductions are still one of the potential pathways of introductions of the species (Hubilla et al. 2007; Santos and Velez-Gavilan, 2024). While invasiveness risk assessment frameworks such as the Aquatic Species Invasiveness Screening Kit (AS-ISK) have identified several high-risk lakes in the country (To et al. 2022; Gilles et al. 2023, 2024, 2025), our predictive models provide the necessary maps for targeted intervention/mitigation to prevent further introduction of this highly invasive fish.

### Model limitations and recommendations

While the study provides a robust framework for predicting *P. disjunctivus* distribution in the Philippines, we acknowledge several limitations. First, the model was developed using a relatively small set of points (n = 11). Although we addressed this using bootstrap subsampling approach and the utilization of SEDI as the validation test - both of which are specifically designed for data-deficient and rare species-the model may not capture the realized niche of the species across every micro-climate in the country, compared to using more occurrences.

Second, our predictors were limited to climate, hydrology, and topography (BIO10, Slope, and Flow accumulation), when in reality, their distribution is also influenced by water quality parameters such as dissolved oxygen, turbidity, pH levels, and salinity (Kumar et al. 2018; Suresh et al. 2019; Santos and Velez-Gavilan, 2024), all of which are not freely available in the country. Hence, adding these important parameters may significantly improve predictions.

Lastly, as with other species distribution models, our maps represent habitat suitability rather than confirmed presence. The actual expansion and invasion of the species remain heavily dependent on human-mediated introductions (Santos and Velez-Gavilan, 2024). Future research should integrate the projected maps with field surveys to truly confirm and mitigate future introduction and invasion of the species.

## CONCLUSION

Using a robust MaxEnt model, we have predicted the species distribution of the highly invasive *Pterygoplichthys disjunctivus* in the Philippines under different socio-economic pathways. By using a representative predictor for climate (BIO10), hydrology (flow accumulation), and topography (slope), our study reveals that the environmental niche of *P. disjunctivus* in the country is primarily constrained by physical landscape features rather than volatile climatic shifts. Given the stability of the species’ habitat suitability across both pessimistic and optimistic scenarios, our findings underscore the urgent need for targeted basin and river system management. Our maps provide strategic maps to implement early-detection and surveillance protocols, ensuring the long-term protection of the Philippines’ freshwater ecosystems.

## STATEMENTS AND DECLARATIONS

The authors have no competing interests to declare that are relevant to the content of this article.

## Data availability

The data from this study is available upon a reasonable request

