## Supplementary figures for "Persistent Invasion Risk: Modeling the near-Current and Future Distribution of Pterygoplichthys disjunctivus across the Philippine Archipelago"

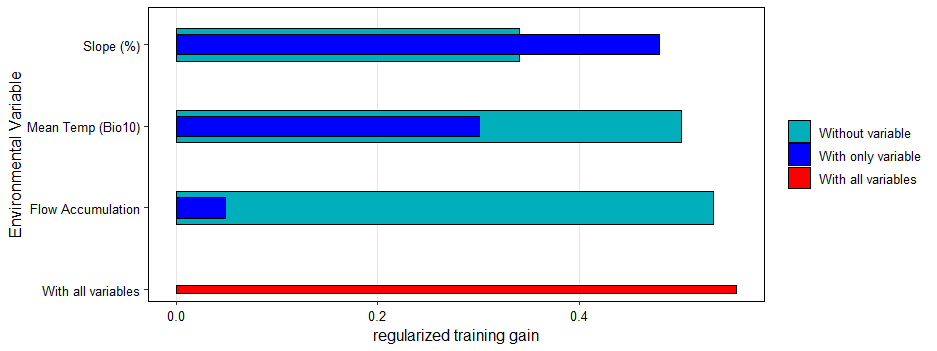

**Figure 1. Jackknife of predictor importance from the MaxEnt model of *P. disjunctivus* using 10 replicates of bootstrap subsampling approach**

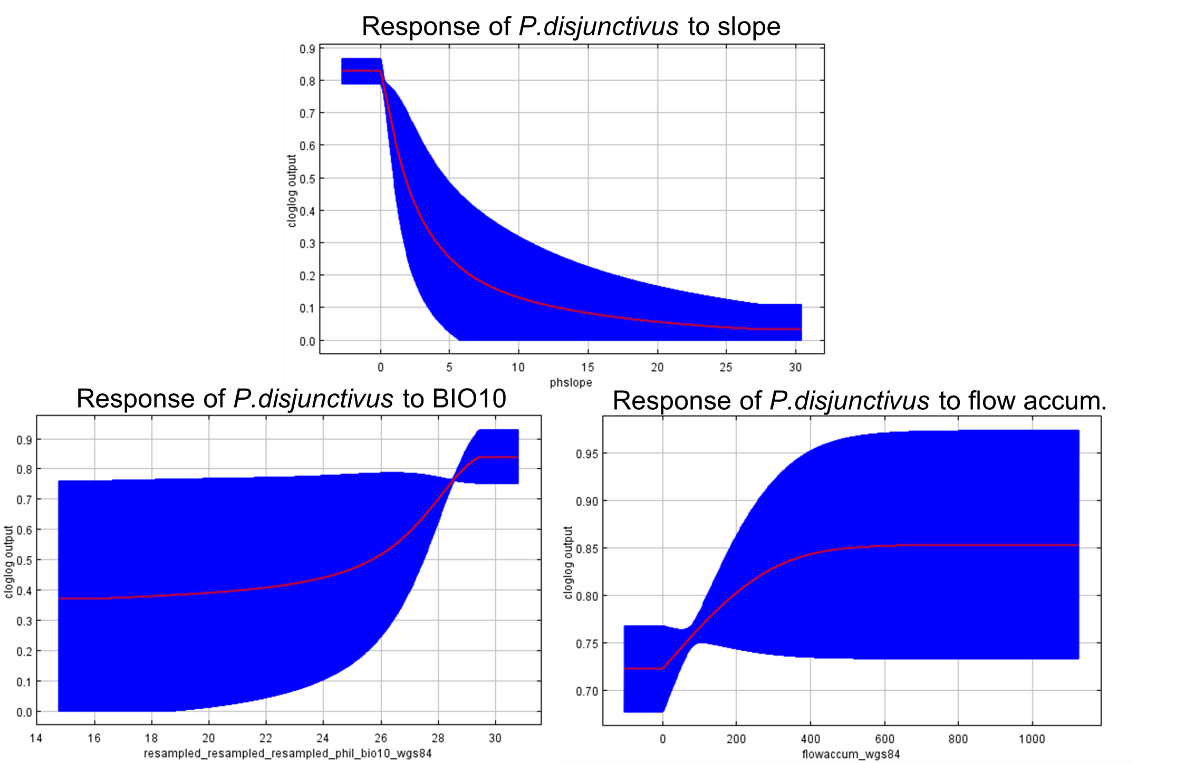

**Figure 2. Response curves of *P. disjunctivus* to the three environmental predictors**
